# A Combinatorial Approach for Single-cell Variant Detection via Phylogenetic Inference

**DOI:** 10.1101/693960

**Authors:** Mohammadamin Edrisi, Hamim Zafar, Luay Nakhleh

## Abstract

Single-cell sequencing provides a powerful approach for elucidating intratumor heterogeneity by resolving cell-to-cell variability. However, it also poses additional challenges including elevated error rates, allelic dropout and non-uniform coverage. A recently introduced single-cell-specific mutation detection algorithm leverages the evolutionary relationship between cells for denoising the data. However, due to its probabilistic nature, this method does not scale well with the number of cells. Here, we develop a novel combinatorial approach for utilizing the genealogical relationship of cells in detecting mutations from noisy single-cell sequencing data. Our method, called scVILP, jointly detects mutations in individual cells and reconstructs a perfect phylogeny among these cells. We employ a novel Integer Linear Program algorithm for deterministically and efficiently solving the joint inference problem. We show that scVILP achieves similar or better accuracy but significantly better runtime over existing methods on simulated data. We also applied scVILP to an empirical human cancer dataset from a high grade serous ovarian cancer patient.

## 1 Introduction

Recently developed single-cell sequencing technologies provide unprecedented resolution in studying genetic variability within a complex tissue [27, 19]. By enabling the sequencing of the genomes of individual cells, these approaches help in delineating the genomic landscape of rare cell populations and provide insights into the somatic evolutionary process. While having single-cell resolution can have major impact on our understanding in neurobiology, immunobiology and other domains [27], it is particularly useful for investigating intratumor heterogeneity and clonal evolution in cancer [20, 29]. Interplay of mutation and selection gives rise to intratumor heterogeneity [21, 18], which contributes to tumor progression and metastasis [25] and can cause tumor relapse via drug resistance [10, 2]. To-date, tumors were mostly analyzed by sequencing bulk samples that consist of admixture of DNA from thousands to millions of cells [9, 22]. Such datasets provide aggregate variant allele frequency (VAF) profiles of somatic mutations, which are subjected to further deconvolution into tumor subpopulations [6]. However, VAFs are noisy, provide limited resolution in identifying rare subpopulations [19] and their deconvolution into clones depends on strong assumptions and therefore may not hold in practice [1].

By enabling the direct measurement of cellular genotypes, single-cell sequencing (SCS) data resolves intratumor heterogeneity to a single-cell level [8, 14]. However, the preparation of SCS data calls for whole genome amplification (WGA) process that can amplify the limited DNA material of a single cell to meet the amount of DNA needed for sequencing [29]. This extensive genome amplification elevates the noise level in SCS data and creates a unique error profile different from bulk sequencing [29]. Technical errors in SCS data include nonuniform coverage, allelic dropout (ADO), false-positive (FP) and false-negative (FN) errors [19, 29]. ADO, i.e., preferential nonamplification of one allele in a heterozygous mutation can result in the loss of the mutated allele removing all evidences of the mutation. On the other hand, random errors introduced during the early stages of amplification can appear as false positive artifacts due to uneven allelic amplifications. The nonuniform coverage in SCS further results in missing data at the sites with insufficient coverage.

Since traditional next-generation sequencing (NGS) variant callers such as GATK [5] do not account for SCS specific errors, variant callers such as Monovar [31] and SCcaller [7] have been specifically designed to deal with the technical errors in SCS. Monovar pools sequencing data across cells to address the problem of low coverage and probabilistically accounts for ADO and FP errors. SCcaller independently detects variants in each cell and accounts for local allelic amplification biases. In another direction, tumor phylogeny inference methods [12, 30, 28] that account for the errors have been developed for studying tumor heterogeneity. A new variant caller SCIΦ [23] combines single-cell variant calling with the reconstruction of the tumor phylogeny. It leverages the fact that somatic cells are related by a phylogeny and employs a Markov Chain Monte Carlo algorithm for calling mutations in each single cell. However, being a probabilistic method, SCIΦ does not scale well with the size of the data and takes a long time to converge on a solution. At the same time, depending on the starting point, it might converge to different solutions.

Here, we present a novel combinatorial formulation for jointly identifying mutations in single cell data by employing the underlying phylogeny. Our approach, scVILP (**s**ingle-**c**ell **V**ariant calling via **I**nteger **L**inear **P**rogram) assumes that the somatic cells evolve along a phylogenetic tree and mutations are acquired along the branches following the infinite sites model as have been used in previous bulk and single-cell studies [6, 12, 23]. We aim to identify the set of single-nucleotide variants in the single cells and genotype them in such a way so that it maximizes the probability of the observed read counts and also the cells are placed at the leaves of a perfect phylogeny that satisfies the infinite sites assumption (ISA). Our solution is deterministic and we solve this problem using a novel Integer Linear Program (ILP) that achieves similar accuracy as SCIΦ but performs significantly better than SCIΦ in terms of runtime. For whole-exome sequencing (WES) data consisting of large numbers of somatic mutation loci, running ILP-solvers can result in high memory consumption. To address this, we introduce a divide-and-conquer version of scVILP where the data matrix is partitioned by columns, scVILP is run on the subsets, and then the results are merged via another ILP formulation to resolve any violations of the ISA assumption in the full dataset. The supertree-based approach achieves similar accuracy as SCIΦ but is much faster.

## 2 Method

In this section, we formulate the joint inference of mutations in single cells and the cellular phylogeny as a combinatorial optimization problem. Following [23], we start by first identifying potential mutation loci and then solve the joint inference problem using a novel Integer Linear Program (ILP).

### 2.1 Input Data

The input data for scVILP corresponds to a read count matrix for *n* cells in the dataset. Let us assume that we identify *m* loci as putative mutated loci. For the joint inference, our input is an *n* × *m* matrix, *X*_*n*×*m*_ = (*X*_*ij*_), where *X*_*ij*_ = (*r*_*ij*_, *v*_*ij*_) denotes the reference and variant read counts for cell *i* and locus *j*. *r*_*ij*_ + *v*_*ij*_ is the coverage at this locus.

### 2.2 Model for Amplification Error

WGA process introduces false positive and false negative errors in single-cell data. We assume that *α* and *β* respectively denote false positive and false negative error rates of single-cell data. We introduce a binary variable *Y*_*ij*_ that denotes the true genotype of mutation *j* in cell *i*. *Y*_*ij*_ = 1 and *Y*_*ij*_ = 0 respectively denote the presence and absence of the mutation. We introduce another variable *D*_*ij*_ that denotes the amplified genotype (after introduction of WGA errors) of mutation *j* in cell *i*. Following previous WGA error models [12, 30], we introduce the following likelihood scheme:

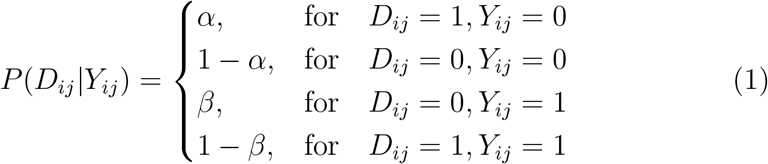

The above likelihood scheme (Eq. (1)) can also be rewritten as:

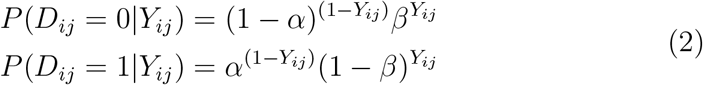

### 2.3 Read Count Model

The read counts (*r*_*ij*_, *v*_*ij*_) at locus *j* of cell *i* is modeled using a Beta-Binomial distribution as commonly used for bulk sequencing data [9] given by:

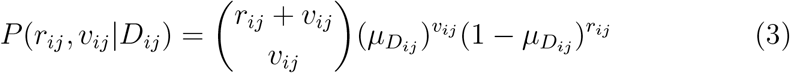

where, 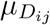 denotes the parameter of the Binomial distribution and it represents the probability of drawing a variant read. This parameter depends on the amplified genotype and is a Beta distributed variable. When *D*_*ij*_ = 0, 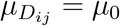 denotes the probability of observing a variant read due to sequencing error, for *D*_*ij*_ = 1, 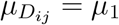 denotes the probability of observing a variant read. Given the amplified genotype *D*_*ij*_, the probability of observing read count (*r*_*ij*_, *v*_*ij*_) is given by

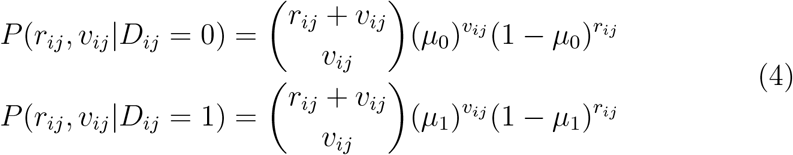

The joint probability of read counts and amplified genotype given the true genotype can be computed by multiplying the probabilities in Eq. (2) and Eq. (4) as given by

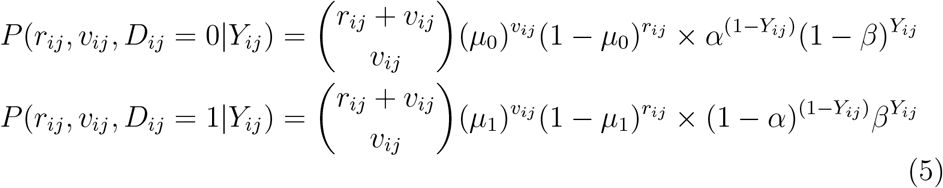

### 2.4 Likelihood Function

The likelihood function of the true genotypes for the cells for an *n* × *m* read count matrix is given by

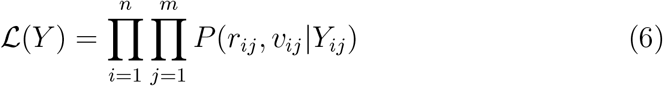

For computing *P* (*r*_*ij*_, *v*_*ij*_|*Y*_*ij*_), we treat the amplified genotype *D*_*ij*_ as a nuisance parameter and marginalize it out as given below:

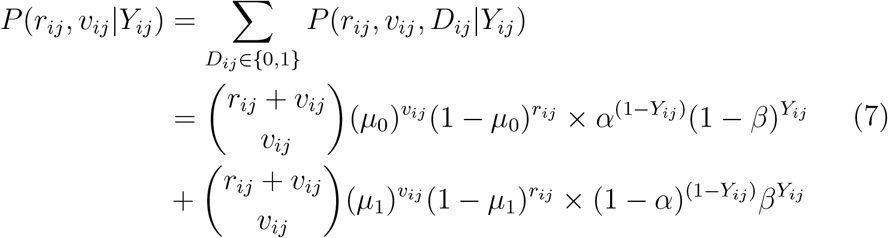

If we denote the Binomial distributions as below:

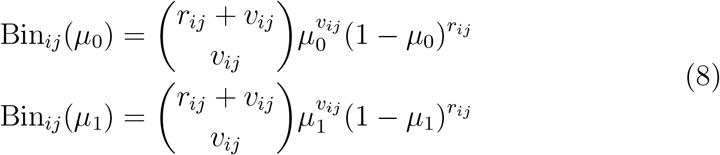

then *P* (*r*_*ij*_, *v*_*ij*_|*Y*_*ij*_) can be rewritten as:

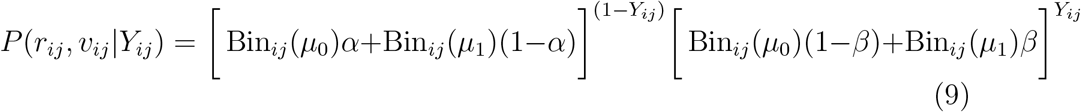

Here our goal is to find a matrix *Y* such that the likelihood function defined in Eq. (6) is maximized. Taking the logarithm of the likelihood function in Eq. (6) and negating it, we obtain the negative log-likelihood function:

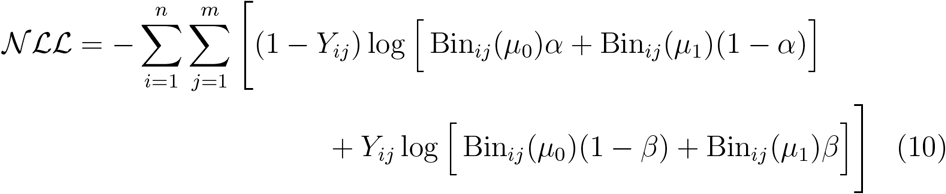

Since this function is linear with respect to the variables *Y*_*ij*_, we can solve this by providing it as the objective function to an ILP solver.

### 2.5 ILP Constraints Pertaining to Perfect Phylogeny

We represent the tumor phylogeny as a perfect phylogeny (PP), leaves of which represent the sampled *n* cells. A perfect phylogeny satisfies the “infinite sites assumption” (ISA), which states that each genomic locus in the dataset mutates at most once during the evolutionary history [15]. We aim to find a genotype matrix *Y* that provides a perfect phylogeny. In order for a binary matrix to be a PP, the three-gametes rule has to hold, i.e., for any given pair of columns (mutations) there must be no three rows (cells) with configuration (1, 0), (0, 1) and (1, 1). 0 and 1 represent the ancestral (unmutated) and mutated state respectively.

In order to enforce that matrix *Y* satisfies the perfect phylogeny criterion, we follow the ILP formulation introduced by Gusfield et al. [11]. which introduces variables *B*(*p, q, a, b*) ∈ {0, 1} for each pair of mutation loci *p* and *q* and for each (*a, b*) ∈ {(0, 1), (1, 0), (1, 1)}. For each cell *i*, the following constraints are introduced on the variables *B*(*p, q, a, b*), *Y*_*ip*_, and *Y*_*iq*_, *i* ∈ {1, …, *n*} and *p, q* ∈ {1, … , *m*}:

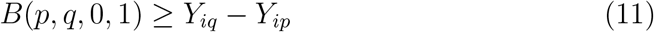

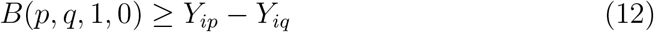

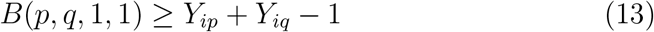

To enforce a perfect phylogeny, the following constraint must hold for all pairs of *p* and *q*:

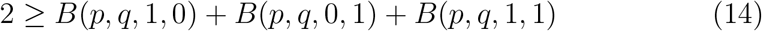

To maximize the likelihood, we need to minimize the negative log-likelihood function defined in Eq. (10), subject to the constraints in Eqns. (11), (12), (13), and (14).

Our ILP formulation can be routinely solved through commercial tools such as CPLEX or Gurobi. In our analysis, we have used Gurobi that concurrently runs multiple deterministic solvers (e.g., dual simplex, primal simplex, etc.) on multiple threads and returns the one that finishes first. As a result, for the same input data and parameter values, Gurobi returns a deterministic solution. It is important to highlight, though, that there could be multiple optimal solutions.

### 2.6 Handling Missing Data

For the loci for which the read count values are missing, we do not consider the related *Y*_*ij*_ variables in the objective function. The values of *Y*_*ij*_ for such missing entries are imputed according to the other values and subject to the Perfect Phylogeny constraints.

### 2.7 Perfect-Phylogeny Supertree

Running ILP-solvers is expensive in terms of memory consumption, specially when the number of mutation loci is high. To overcome this limitation, we use a divide-and-conquer approach consisting of the following steps:

1. Partition the columns of *X*_*n*×*m*_, into *k* subsets of mutation loci, {*C*_1_, · · ·, *C*_*k*_}, where each *C*_*j*_ is a set of columns in *X*_*n*×*m*_. We define the submatrices {*X*^(1)^, · · · , *X*^(*k*)^} as follows:

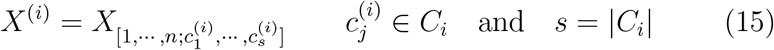
2. For each submatrix, *X*^(*i*)^, optimize the objective function in Eq. 10 (subject to the constraints in Eqns 11, 12, 13, and 14) independently. This step results in genotyped submatrices, {*G*^(1)^, · · ·, *G*^(*k*)^}.
3. Concatenate the genotyped submatrices into *G*_*n*×*m*_ as follows:

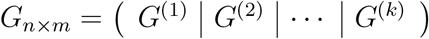
4. If *G*_*n*×*m*_ does not admit a Perfect Phylogeny, find the minimum number of changes to apply on the entries of *G*_*n*×*m*_ such that the modified matrix satisfies the Perfect Phylogeny model. We name this final step as PERFECT PHYLOGENY CORRECTION PROBLEM (PPC), which is described in the following section.

### 2.8 The Perfect Phylogeny Correction Problem

We define the PERFECT PHYLOGENY CORRECTION PROBLEM as inferring matrix 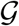 from matrix *G*, which does not satisfy perfect phylogeny such that 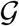 admits a perfect phylogeny. PPC consists of two steps:

1. Find the minimum number of characters (mutation loci) which must be removed in order to make the matrix, *G* admit a perfect phylogeny. This problem is known as CHARACTER-REMOVAL PROBLEM (CRP) [11].
2. Among the entries of the selected loci in the previous step, minimize the number of changes to apply such that final matrix 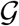 is perfect phylogeny. We name this problem as ENTRY-CHANGE PROBLEM (ECP).

#### 2.8.1 The Character-Removal Problem

Given a genotyped matrix *G*, which does not admit a PP, we aim to find the smallest set of mutation loci to remove such that the final result admits a perfect phylogeny. The three-gametes rule is violated by the pairs of columns (loci) whose corresponding rows (cells) contain all three configurations (0, 1), (1, 0), and (1, 1). As Gusfield *et al.* [11] showed, there is an efficient ILP formulation for solving this problem.

Let *δ*(*i*) be a binary variable used to indicate whether or not loci *i* will be removed. Then for each pair of mutation loci (*p, q*), *p, q* ∈ {1, · · ·, *m*} which violate three-gametes rule, we add the inequality *δ*(*p*) + *δ*(*q*) ≥ 1. Finally, we minimize the objective function 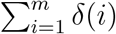. An example of CRP is shown in Table 1.

**Table 1:**
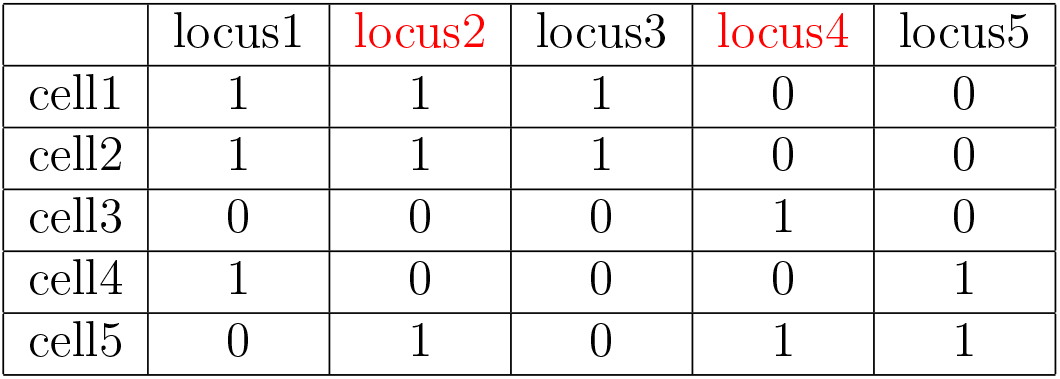
Character-Removal algorithm selects two loci, 2 and 4 to remove from *G*.

|  | locus1 | locus2 | locus3 | locus4 | locus5 |
| --- | --- | --- | --- | --- | --- |
| cell1 | 1 | 1 | 1 | 0 | 0 |
| cell2 | 1 | 1 | 1 | 0 | 0 |
| cell3 | 0 | 0 | 0 | 1 | 0 |
| cell4 | 1 | 0 | 0 | 0 | 1 |
| cell5 | 0 | 1 | 0 | 1 | 1 |

#### 2.8.2 The Entry-Change Problem

We define the ENTRY-CHANGE PROBLEM as inferring a matrix 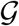 from *G* by changing the entries of *G* such that 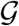 is a PP. Here, the character-removal problem plays an important role by returning the set of loci to be removed from *G* in order to admit a PP. As a result, instead of the whole matrix *G*, ECP can be applied to this set of loci. This in turn reduces the search space for solving ECP. Let *R* = {*r*_1_, · · ·, *r*_*l*_} denote this set of mutations.

The ILP for ECP introduces variables *Change*(*i, j*) ∈ {0, 1} for each cell *i* ∈ {1, · · ·, *n*} and *j* ∈ *R* indicating whether or not the entry *G*(*i, j*) will be changed. We also use the variables *B*(*p, q, a, b*) ∈ {0, 1} described in section 2.5 with the same defintion. The program next creates inequalities that force the binary variable *B*(*p, q, a, b*) to 1 if changing (or not changing) either or both entries *G*(*i, p*) and *G*(*i, q*) produces the combination (*a, b*). To explain the formulation in detail, consider the pair of columns shown in Table 2. The first row creates the combination (1,1) between locus 1 and 2 if the entry *G*(1, 2) does not change. This condition can be formulated as the following inequality:

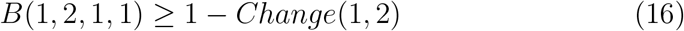

**Table 2:** Five rows and two columns in the input *G* of ECP.

|  | locus1 | locus2 |
| --- | --- | --- |
| cell1 | 1 | 1 |
| cell2 | 1 | 1 |
| cell3 | 0 | 0 |
| cell4 | 1 | 0 |
| cell5 | 0 | 1 |

And chaning the entry *G*(1, 2) creates the combination (1, 0):

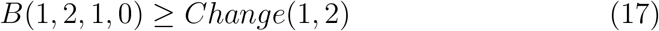

The constraints are built similarly, according to the other rows and columns. The constraint on variables *B*(*p, q, a, b*) which guarantees PP is the same as the inequality 14. The objective function is to Minimize 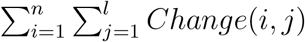 where *n* is the number of cells and *l* is the number of problematic loci.

While [3] presented an ILP solution for finding a minimum flip supertree admitting a PP, our combination of character removal and entry change problem does not guarantee a minimum flip supertree. Here our goal is to find a Perfect phylogeny supertree and our two-step approach guarantees to achieve that and at the same time restricting ECP to the sites selected by character removal results in reduction of runtime.

## 3 Results

### 3.1 Benchmarking on Simulated Data

To benchmark the performance of our approach and compare it with that of SCIΦ we first ran both of the methods on simulated datasets for which the ground truth is known. For generating simulated datasets, we used the single-cell read count simulator module introduced in [23]. The simulator first generates a model tumor tree, which starts from a progenitor cell without any mutation. Cells evolve along the branches of the tree and the mutations are acquired along these branches. The leaves of the tree represent the single cells. For simulating allelic dropout, mutations present in the cells at the leaves of the phylogeny are dropped out randomly with probability *β* (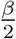 of the mutations become wild type and 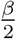 become homozygous). This simulator mimics the noisy MDA process [4] and accounts for the sequencing error whose corresponding probability values were set to 5 × 10^−7^ and 10^−3^ respectively. The MDA process was repeated 50 times as suggested by the authors of SCIΦ [23]. The mean and variance associated with the coverage distribution were set to 25 and 50 respectively. Finally, we also introduced 10% missing data.

We investigated how each method performs on datasets carrying different number of cells. The number of cells in the datasets, *n*, was varied as *n* = 50, and *n* = 100. The number of sites, *m*, was set at *m* = 100. We generated 5 datasets for each value of *n*. The rate of allelic dropout was chosen from the values {0.05, 0.1, 0.15, 0.2, 0.25}. SCIΦ was run according to the authors’ recommendation with an additional filter of requiring at least two cells to show variant read count of at least three. Our method scVILP was run with allelic dropout values given as input parameters to the program. In order to emulate the real settings where we do not know the exact values of the allelic dropouts, we added noise to their true values in the same way described in [16]. To add noise to the dropout values used as input to scVILP, a random value was sampled from a normal distribution with mean 1 and standards deviation 0.1 from the interval (0.5, 2) and multiplied with the dropout values. The parameter *μ*_0_ was empirically determined (by varying between 10^−8^ − 10^−3^) and set to 10^−5^. The value of the parameter *μ*_1_ was set to 0.5 assuming heterozygous somatic mutation. The results are shown in Fig. 1. Performance of each method was measured in terms of precision and recall. Precision is defined as 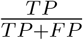, whereas recall is defined as 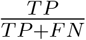 (TP: true positive, FP: false positive and FN: false negative). For each setting, scVILP achieved similar or better precision than SCIΦ. On the other hand, SCIΦ achieved a little better recall than scVILP for *n* = 100, but scVILP’s recall was comparable to that of SCIΦ for *n* = 50. Overall, both methods achieved similar precision and recall. This can be observed when comparing the F1 measure, which is the harmonic mean of precision and recall. However, scVILP’s median F1 score was marginally better than that of SCIΦ for each setting. These experiments show that both of these methods achieve similar accuracy in detecting the mutations. Since both of these methods utilize the underlying phylogenetic tree (assuming infinite sites assumption) in inferring the variants, both of them can identify mutations in a particular cell even if there is very little or missing variant support at a specific locus.

**Figure 1:**
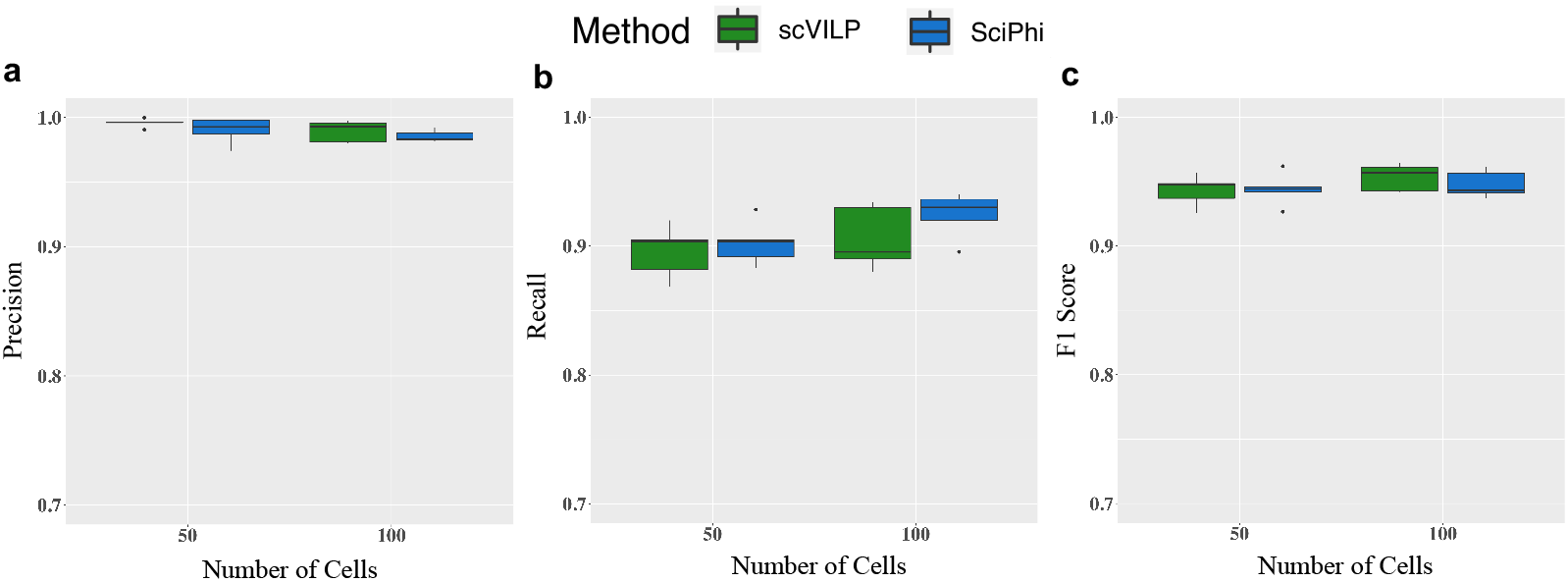
Performance of scVILP and SCIΦ on simulated data with different number of cells. The number of mutations is 100. Performance measured as (a) precision, (b) recall, and (c) F1 score.

However, the major difference between these two methods lie in the use of the inference algorithm. While SCIΦ uses a Markov Chain Monte Carlo algorithm for sampling from the posterior distribution, scVILP solves the problem deterministically using ILP. Consequently, scVILP is much faster in finding the solution as can be seen in the runtime comparison (Fig. 2). For *n* = 50, scVILP was on average ~ 38 times faster than SCIΦ, while for *n* = 100, it was on average ~ 31 times faster than SCIΦ.

**Figure 2:**
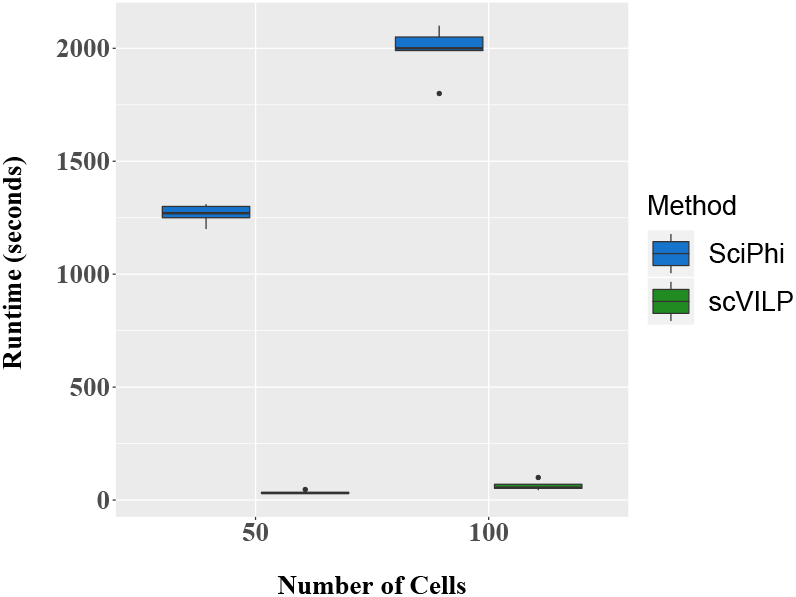
Runtime comparison for scVILP and SCIΦ on simulated data with different number of cells. The number of mutations is 100.

Allelic dropout is a major source of error in single-cell data [19]. We further assessed scVILP’s performance under different values of allelic dropout rate. We generated 5 datasets for each value of allelic dropout in {0.2, 0.3}. For these datasets, we used *n* = 50 and *m* = 100. Both scVILP and SCIΦ were compared on these datasets in terms of precision and recall (Fig. 3). The F1 score for each method decreased with an increase in the allelic dropout rate. For each setting of allelic dropout rate, scVILP had slightly higher precision than that of SCIΦ. Recalls for both methods were comparable; however, scVILP had slightly better recall for the datasets with higher allelic dropout rate. This experiment demonstrates that scVILP performs robustly for varying allelic dropout rate.

**Figure 3:**
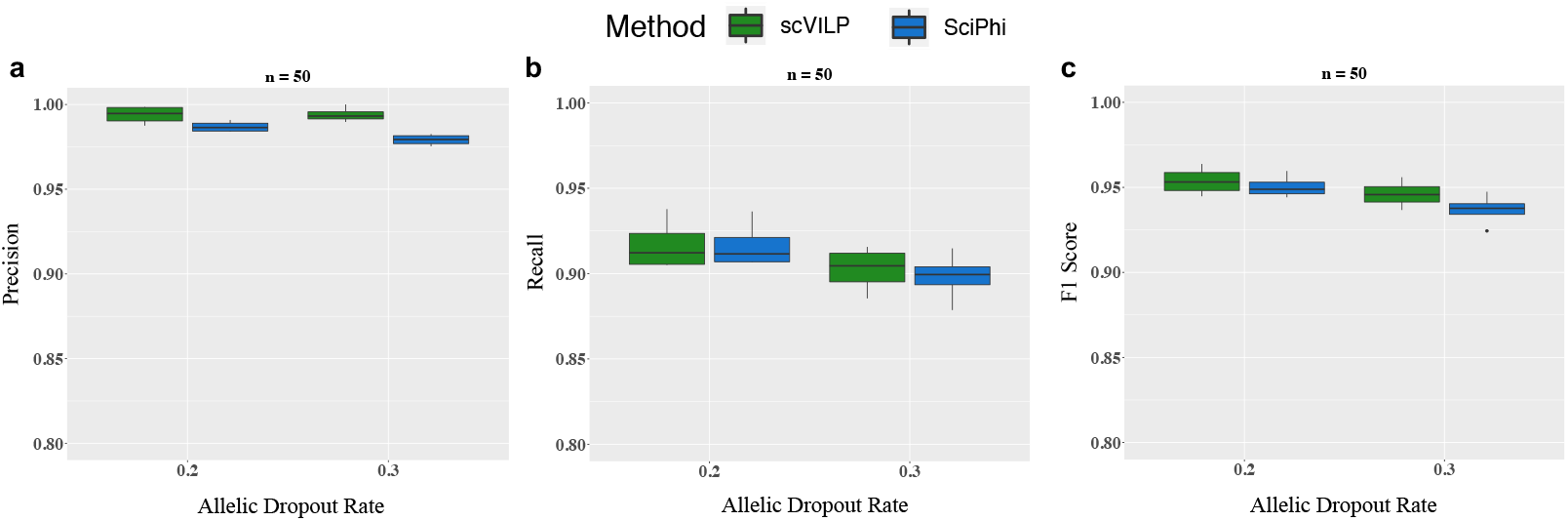
Performance of scVILP and SCIΦ on simulated data with different allelic dropout rates. The number of mutations is 100 and number of cells is 50. Performance measured as (a) precision, (b) recall, and (c) F1 score.

Due to the high memory consumption of ILP-solvers, straightforward application of scVILP might run into memory issues for very large number of mutations that can occur in single-cell WES datasets. For addressing such cases, we developed the Perfect Phylogeny supertree algorithm that should run without any memory issue, reducing the running time while maintaining tolerable tradeoff in accuracy. To analyze its performance, we generated three datasets containing 100 cells and 1000 mutation loci with allelic dropout chosen from {0, 0.15, 0.25}. We compared scVILP against SCIΦ on these datasets (Fig. 4). While scVILP and SCIΦ had similar F1 score for these datasets, scVILP had significantly shorter runtime compared to SCIΦ. For these datasets, direct ILP formulation of scVILP failed to run on a machine with 32GB RAM, but the Perfect Phylogeny supertree algorithm, by dividing the original matrix into 5 submatrices, ran successfully with 11GB peak memory usage. Since SCS technologies are generating SNV data on hundreds of cells, but not yet on thousands of cells, we did not investigate the performance of scVILP and SCIΦ by simulating larger number of cells.

**Figure 4:**
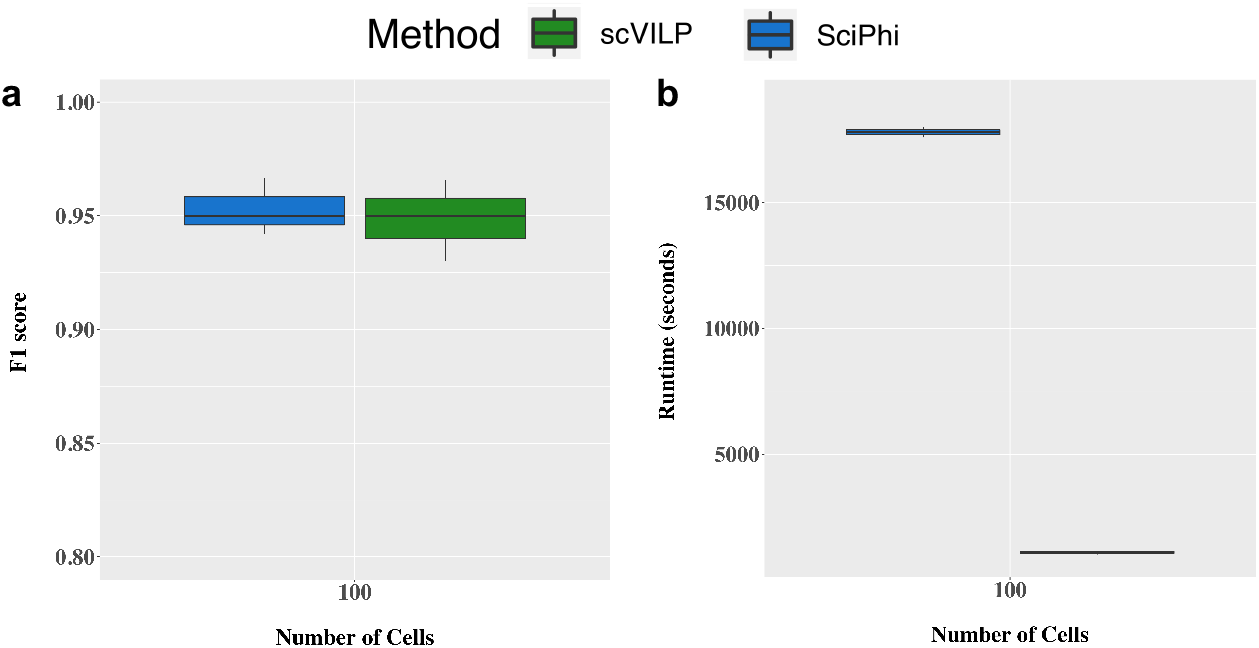
Performance of scVILP’s Perfect Phylogeny supertree algorithm compared to SCIΦ on simulated data with large number of mutations. The number of mutations is 1000 and the number of cells is 100. (a) comparison of F1 score, (b) runtime comparison.

### 3.2 Application of scVILP on Real Data

Finally, we applied scVILP on a human cancer dataset. This dataset consists of 370 single cells and 43 mutation loci from a high-grade serous ovarian cancer patient described in [17]. For comparison, we also ran SCIΦ on this dataset and the mutation calls for both methods are shown in Fig. 5. Mutation calls for both these methods were mostly similar. The underlying phylogenetic trees inferred by these methods were also similar (Robinson-Foulds distance 0.123, computed by PAUP[24]) indicating that multiple perfect phylogenies fit the data. We further analyzed the mutation calls that were different between the methods. A large number of mutations that were genotyped as 1 by SCIΦ and 0 by scVILP had high reference read count and low variant read count indicating scVILP’s genotyping might have been more accurate for these sites. Both methods were run on a PC with 8GB RAM and 2 GHz Intel Core i5 processor. scVILP significantly outperformed SCIΦ in terms of runtime. The runtime for scVILP was around 30 minutes compared to the nearly 12 hours runtime of SCIΦ.

**Figure 5:**
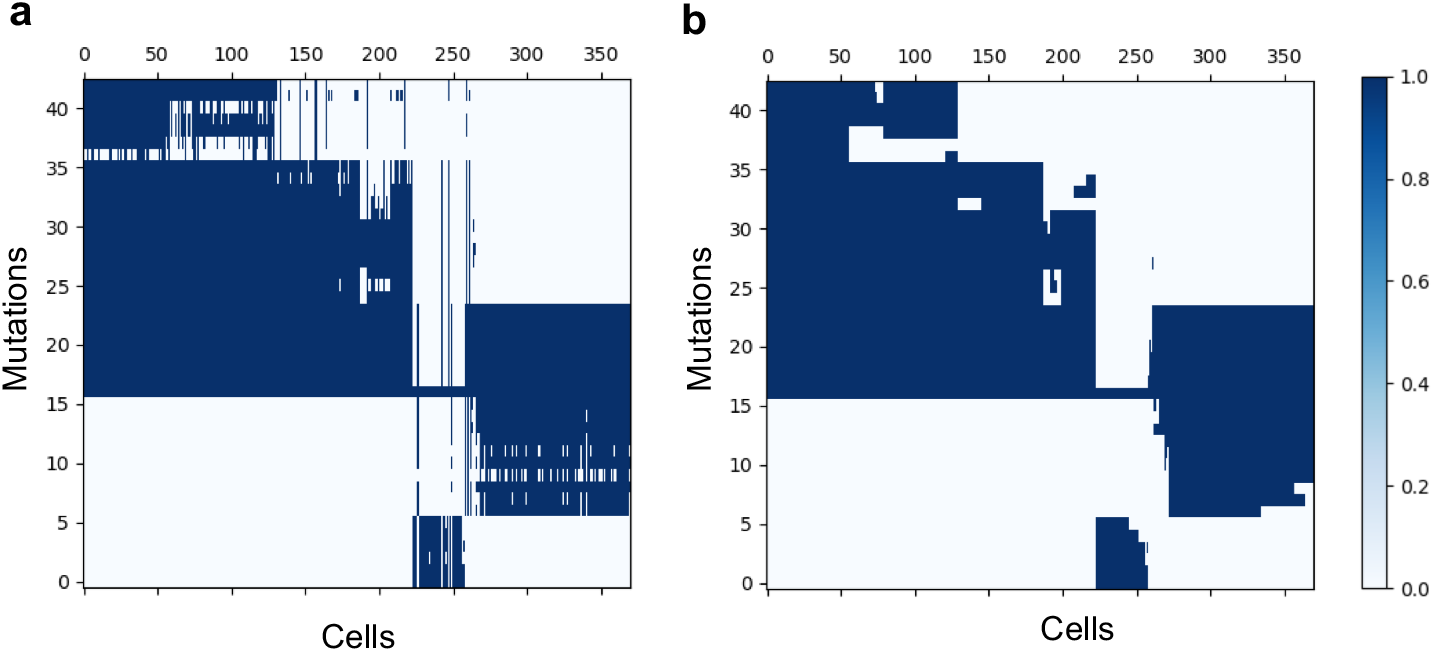
Summary of the mutation calls from SCIΦ and scVILP on a dataset consisting of 370 cells from a patient with high-grade serous ovarian cancer [17]. Deep blue and white entries in the heatmaps represent mutations states 1 and 0, respectively. (a) Mutation calls of SCIΦ. (b) Mutation calls of scVILP.

## 4 Conclusions

Here, we introduced scVILP, a novel combinatorial approach for jointly inferring the mutations in single cells and a perfect phylogeny that connects the cells. Using a novel Integer Linear Program, our method infers the mutations in individual cells assuming they evolve along a perfect phylogeny. scVILP probabilistically models the observed read counts but combinatorially solves the joint inference problem.

We compared scVILP to SCIΦ [23] on both simulated and real datasets. For the simulated data, both methods performed similarly in terms of precision and recall (as shown by their F1 scores). However, scVILP significantly outperformed SCIΦ in terms of runtime. Similarly for real human cancer dataset, scVILP achieved a solution much faster compared to SCIΦ. For the real dataset, SCIΦ produced different solutions in different runs because of its probabilistic nature. For the real dataset, even though mutation calls from scVILP and SCIΦ were similar, we also observed certain differences indicating that our deterministic algorithm and SCIΦ’s probabilistic inference do not necessarily arrive at a unique solution in spite of the use of underlying perfect phylogenetic tree. This also indicates that potentially, multiple perfect phylogenies can fit the data well necessitating the development of methods that also assess uncertainty in the inferences.

Our method could potentially be improved by including violations of the perfect phylogeny. Tumor phylogeny studies [30, 13, 26] show possible violations of ISA in single-cell sequencing datasets. More comprehensive evaluation of these violations could be performed by analyzing the read counts in conjunction with tumor phylogeny inference. Another improvement would be the inclusion of copy number data.

As single-cell sequencing becomes more high-throughput producing thousands of cells, scVILP will be very helpful for identifying the mutations from such larger datasets. scVILP is very fast and accurate in detecting the mutations as well as the phylogeny underlying the cells. Addressing both these problems in a single combinatorial framework will be very helpful for quick analysis of large datasets for understanding tumor heterogeneity.

## References

[1] Niko Beerenwinkel, Roland F Schwarz, Moritz Gerstung, and Florian Markowetz. Cancer evolution: mathematical models and computational inference. Systematic Biology, 64(1):e1–e25, 2014.

[2] Rebecca A Burrell and Charles Swanton. Tumour heterogeneity and the evolution of polyclonal drug resistance. Molecular Oncology, 8(6):1095–1111, 2014.

[3] Markus Chimani, Sven Rahmann, and Sebastian Böcker. Exact ILP solutions for phylogenetic minimum flip problems. In Proceedings of the First ACM International Conference on Bioinformatics and Computational Biology, pages 147–153. ACM, 2010.

[4] Frank B Dean, John R Nelson, Theresa L Giesler, and Roger S Lasken. Rapid amplification of plasmid and phage DNA using phi29 DNA polymerase and multiply-primed rolling circle amplification. Genome Research, 11(6):1095–1099, 2001.

[5] Mark A. DePristo, Eric Banks, Ryan Poplin, Kiran V. Garimella, Jared R. Maguire, Christopher Hartl, Anthony A. Philippakis, Guillermo del Angel, Manuel A. Rivas, Matt Hanna, Aaron McKenna, Tim J. Fennell, Andrew M. Kernytsky, Andrey Y. Sivachenko, Kristian Cibulskis, Stacey B. Gabriel, David Altshuler, and Mark J. Daly. A framework for variation discovery and genotyping using next-generation DNA sequencing data. Nature Genetics, 43(5):491–498, 2011.

[6] Amit G. Deshwar, Shankar Vembu, Christina K. Yung, Gun Ho Jang, Lincoln Stein, and Quaid Morris. Phylowgs: Reconstructing subclonal composition and evolution from whole-genome sequencing of tumors. Genome Biology, 16(1):1–20, 2015.

[7] Xiao Dong, Lei Zhang, Brandon Milholland, Moonsook Lee, Alexander Y Maslov, Tao Wang, and Jan Vijg. Accurate identification of single-nucleotide variants in whole-genome-amplified single cells. Nature Methods, 14(5):491, 2017.

[8] Charles Gawad, Winston Koh, and Stephen R. Quake. Dissecting the clonal origins of childhood acute lymphoblastic leukemia by single-cell genomics. Proceedings of the National Academy of Sciences, 111(50):17947–17952, 2014.

[9] Moritz Gerstung, Christian Beisel, Markus Rechsteiner, Peter Wild, Peter Schraml, Holger Moch, and Niko Beerenwinkel. Reliable detection of subclonal single-nucleotide variants in tumour cell populations. Nature Communications, 3:811, 2012.

[10] Robert J. Gillies, Daniel Verduzco, and Robert A. Gatenby. Evolutionary dynamics of carcinogenesis and why targeted therapy does not work. Nat Rev Cancer, 12(7):487–493, Jul 2012.

[11] Dan Gusfield, Yelena Frid, and Dan Brown. Integer programming formulations and computations solving phylogenetic and population genetic problems with missing or genotypic data. In International Computing and Combinatorics Conference, pages 51–64. Springer, 2007.

[12] Katharina Jahn, Jack Kuipers, and Niko Beerenwinkel. Tree inference for single-cell data. Genome Biology, 17(1):1–17, 2016.

[13] Jack Kuipers, Katharina Jahn, Benjamin J Raphael, and Niko Beerenwinkel. Single-cell sequencing data reveal widespread recurrence and loss of mutational hits in the life histories of tumors. Genome research, 27(11):1885–1894, 2017.

[14] Marco L Leung, Alexander Davis, Ruli Gao, Anna Casasent, Yong Wang, Emi Sei, Eduardo Vilar, Dipen Maru, Scott Kopetz, and Nicholas E Navin. Single-cell DNA sequencing reveals a late-dissemination model in metastatic colorectal cancer. Genome Research, 27(8):1287–1299, 2017.

[15] Jian Ma, Aakrosh Ratan, Brian J. Raney, Bernard B. Suh, Webb Miller, and David Haussler. The infinite sites model of genome evolution. Proceedings of the National Academy of Sciences, 105(38):14254–14261, 2008.

[16] Salem Malikic, Simone Ciccolella, Farid Rashidi Mehrabadi, Camir Ricketts, Md Khaledur Rahman, Ehsan Haghshenas, Daniel Seidman, Faraz Hach, Iman Hajirasouliha, and S Cenk Sahinalp. PhISCS-a combinatorial approach for sub-perfect tumor phylogeny reconstruction via integrative use of single cell and bulk sequencing data. BioRxiv, page 376996, 2018.

[17] Andrew McPherson, Andrew Roth, Emma Laks, Tehmina Masud, Ali Bashashati, Allen W Zhang, Gavin Ha, Justina Biele, Damian Yap, Adrian Wan, et al. Divergent modes of clonal spread and intraperitoneal mixing in high-grade serous ovarian cancer. Nature Genetics, 48(7):758, 2016.

[18] Lauren M.F. Merlo, John W. Pepper, Brian J. Reid, and Carlo C. Maley. Cancer as an evolutionary and ecological process. Nat Rev Cancer, 6(12):924–935, Dec 2006.

[19] Nicholas Navin. Cancer genomics: one cell at a time. Genome Biology, 15(8):452–465, 2014.

[20] Nicholas E. Navin. The first five years of single-cell cancer genomics and beyond. Genome Research, 25(10):1499–1507, Oct 2015.

[21] PC Nowell. The clonal evolution of tumor cell populations. Science, 194(4260):23–28, 1976.

[22] Andrew Roth, Jaswinder Khattra, Damian Yap, Adrian Wan, Emma Laks, Justina Biele, Gavin Ha, Samuel Aparicio, Alexandre Bouchard-Cote, and Sohrab P. Shah. PyClone: statistical inference of clonal population structure in cancer. Nat Meth, 11(4):396–398, Apr 2014.

[23] Jochen Singer, Jack Kuipers, Katharina Jahn, and Niko Beerenwinkel. Single-cell mutation identification via phylogenetic inference. Nature Communications, 9(1):5144, 2018.

[24] DL Swafford. PAUP*. Phylogenetic analysis using parsimony (* and other methods). Vol. Sinauer Associates, Sunderland, MA, 2002.

[25] Charles Swanton. Intratumor heterogeneity: evolution through space and time. Cancer research, 72(19):4875–4882, 2012.

[26] Lili Wang, Jean Fan, Joshua M Francis, George Georghiou, Sarah Hergert, Shuqiang Li, Rutendo Gambe, Chensheng W Zhou, Chunxiao Yang, Sheng Xiao, et al. Integrated single-cell genetic and transcriptional analysis suggests novel drivers of chronic lymphocytic leukemia. Genome research, 27(8):1300–1311, 2017.

[27] Yong Wang and Nicholas E. Navin. Advances and applications of single-cell sequencing technologies. Molecular Cell, 58(4):598–609, 2015.

[28] Hamim Zafar, Nicholas Navin, Ken Chen, and Luay Nakhleh. SiCloneFit: Bayesian inference of population structure, genotype, and phylogeny of tumor clones from single-cell genome sequencing data. bioRxiv, page 394262, 2018.

[29] Hamim Zafar, Nicholas Navin, Luay Nakhleh, and Ken Chen. Computational approaches for inferring tumor evolution from single-cell genomic data. Current Opinion in Systems Biology, 7:16–25, 2018.

[30] Hamim Zafar, Anthony Tzen, Nicholas Navin, Ken Chen, and Luay Nakhleh. SiFit: inferring tumor trees from single-cell sequencing data under finite-sites models. Genome Biology, 18(1):178, 2017.

[31] Hamim Zafar, Yong Wang, Luay Nakhleh, Nicholas Navin, and Ken Chen. Monovar: single-nucleotide variant detection in single cells. Nature Methods, 13(6):505–507, Jun 2016.

